# A chicken interferon stimulated gene screen reveals limited genes restricting influenza replication

**DOI:** 10.64898/2026.07.24.740132

**Authors:** Alison J. Barkhymer, Jordan T. Becker, Frances K. Shepherd, Clayton K. Mickelson, Lauren M. Pross, Mark Salnikov, Ryan A. Langlois

## Abstract

Avian influenza A virus (IAV) is a major global health threat. Not only does it cause significant economic losses to the agricultural industry, but it also presents a risk of spillover into the human population. However, several obstacles must be overcome for a successful cross-species transmission event to occur. One of these obstacles is the innate immunity of the host. While the innate immune response to IAV in humans has been well-studied, the innate immune response to IAV in birds is not as well understood. Here, we explore the interferon-induced immune response by identifying interferon-stimulated genes (ISGs) in chicken, quail, and duck cells, and find that they are largely species-specific. Using this data, we performed a CRISPR knockout ISG screen against two strains of IAV, the mouse-adapted PR8, and the low pathogenicity avian IAV isolate WF10. We find that for PR8, IFITM3 is the dominant restrictor of viral replication with several additional hits with modest effects, consistent with previous work in mammals. For avian WF10, IFITM3 was also a hit however there were two additional hits absent for PR8, the HSP70 co-chaperone protein BAG5 and deubiquitinase OTUD4. Overall, these findings confirm the divergence of interferon-mediated immunity between species and establish a screening system that can be used to identify ISGs important for viral restriction in avian species.

**IMPORTANCE:** Wild birds, especially waterfowl and shore birds, are the primary natural reservoir for influenza A viruses. These birds can transmit IAV to domestic poultry, where the spillover risk to humans increases. Chickens are the most common species of domestic poultry world-wide and are frequently infected with avian influenza viruses. Despite this, relatively little is known about how their immune system responds to influenza infection. This lack of knowledge creates a barrier to finding ways of preventing or mitigating IAV infection in chicken flocks. In this study, we characterized the interferon-induced immune response to IAV in chickens and found that only a small subset of ISGs were responsible for restricting IAV in chicken cells.

## INTRODUCTION

Influenza A virus (IAV) is an omnipresent zoonotic pathogen endemic in global waterfowl and shorebirds frequently entering agricultural poultry flocks, and sporadically infecting mammalian species. In addition to being seasonally endemic in humans, IAV can also cause pandemics, which have likely occurred sporadically over the last millennia. The first to be molecularly verified occurred in 1918, followed by pandemics caused by zoonotic reassortment in 1957, 1968, and 2009. While humans are constantly exposed to avian influenza viruses, cross-species jumps by these viruses are infrequent and sustained transmission in humans remains rare. This is due to many barriers to cross-species infections.

Influenza viruses need to use host factors to complete their life cycle, including sialic acid for entry and ANP32A for viral polymerase activity, which can be divergent across avians and mammals(1). Antiviral responses also pose a strong barrier, are divergent across species, and drive viral evolution. Viruses, in turn, can shape the genome by exerting selective pressure on a population(2). The major intrinsic antiviral defense mechanism in chordates is driven by interferon (IFN) which results in the induction of hundreds of interferon-stimulated genes (ISGs) which make cells inhospitable to virus replication. These genes are frequently under positive selection(3, 4), and because of this, there is significant divergence in IFN responses across species which can pose a barrier to cross species jumps. For example, divergence and presence/absence of ZAP/KHNYN, MX1, and BTN3A3 create a barrier between avian influenza viruses emerging into mammals(5–8).

Most species encode hundreds of ISGs with a wide range of potential antiviral effects, and determining the ISGs capable of restricting the replication of a specific virus in any given species can be a challenge. For avian IAV, this is made even more difficult by the comparative lack of knowledge on the intrinsic innate immune systems of birds. Forward and reverse genetic screens in humans have identified several interferon stimulated genes that can control influenza infection including IFITM3, MX1, MOV10, PKR, ZAP, DDX21, TRIM22, TREX1, HAX1, OAS, VIPERIN(9–13). Despite avian species serving as the main reservoir of IAV and source of high-pathogenicity strains, few genome-wide or subset screens have been performed in avian systems. Several antiviral genes have been uncovered through testing avian orthologs of known mammalian antiviral genes including IFITM3, IFITM5, and OAS/L(14–17). One strategy to discover antiviral genes has been to perform targeted CRISPR KO screens against mammalian ISG which has been done for HIV-1, SARS-CoV-2, Venezuelan equine encephalitis virus, vesicular stomatitis virus, and *Toxoplasma gondii*(*18–24*). This has led to the discovery of many virus-specific ISGs.

Here we determine the antiviral genes that can control influenza virus infection in Chickens. We used RNA-seq to identify ISGs in chicken, duck, and quail cell lines following conspecific IFNα treatment. Then, we designed a CRISPR library based on these ISGs and those identified from the literature. We screened with guide RNAs targeting 543 protein-coding chicken ISGs in the presence and absence of species-specific IFN treatment to identify antiviral factors. We identified chicken IFITM3 as the major antiviral gene inhibiting IAV replication. Finally, we identified several additional ISGs including some that were strain specific.

## RESULTS

### Avian interferon responses are species-specific

To directly compare the interferon responses across representative avian species we treated chicken, duck, and quail immortalized fibroblasts (DF-1, CCL-141, and QT6 cells, respectively) with 200ng/mL of chicken IFNα, duck IFNα, quail IFNα, or 1000units/mL of universal IFN (chimera of human IFNα A/D). mRNA levels of the ISGs OASL and IFIT5 were measured by qRT-PCR 4 hours post-treatment (hpt). Chicken and duck ISG expression was only substantially stimulated by conspecific IFN (Figure 1A). Chicken ISGs were slightly induced by duck IFN, but the fold change was 3 logs lower than with chicken IFN (Figure 1A). The duck ISGs tested were only induced by duck IFN, with virtually no upregulation with chicken, quail, or universal IFN (Figure 1A). Surprisingly, quail ISGs were upregulated at comparable levels by all three avian IFNs, although not by mammalian universal IFN (Figure 1A). To ensure that the IFNs we were using were not contaminated with something that could induce the expression of ISGs, we also treated human A549 cells which should not be responsive to avian IFNs. As expected, only mammalian universal interferon was able to induce the expression of ISGs in A549 cells (Figure 1A).

**Figure 1.**
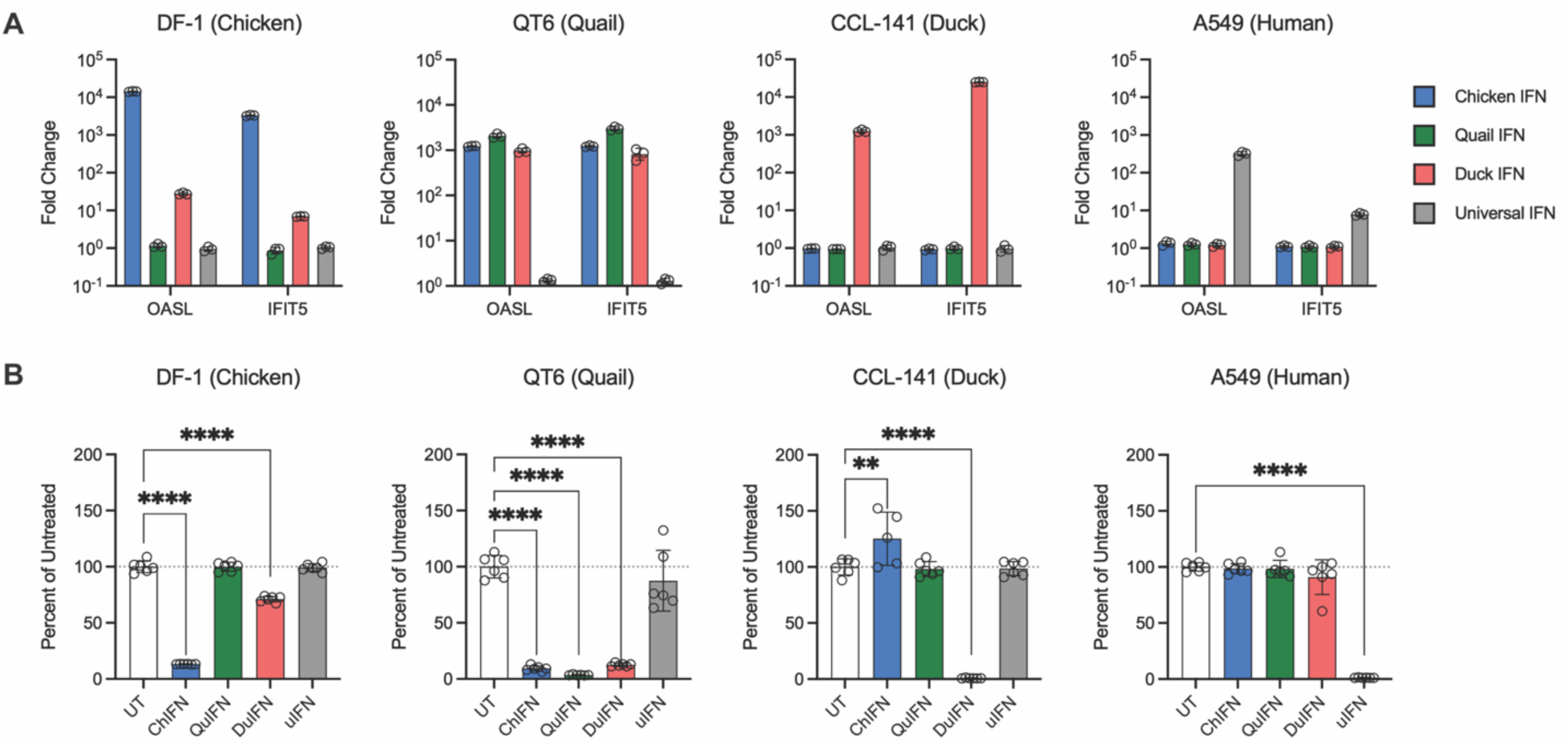
Avian responses to IFNα treatment and PR8 infection are species-specific. **A)** qPCR of ISGs from cells collected 4h post treatment with IFNα from different species. N = 3 samples per group and representative of 2 independent experiments. **B)** Chicken, duck, quail, or human cells were pretreated with IFNα for 24h and infected with scPR8-mCh at MOI 5. Fluorescence measured 24h post infection. Percent infection in untreated cells was normalized to 100, with values for other treatment groups given at a precent of that. N = 6 samples per group, representative of two experiments for avian cells and one experiment for A549.

To test the functionality of the IFNs in the context of a viral infection, each cell line was infected with a single-cycle PR8 influenza A virus that expresses mCherry in place of HA (scPR8-mCh). This allows us to use mCherry fluorescence as a proxy for viral replication(25). Cells were infected with scPR8-mCh at an MOI of 5 following 24h of pretreatment with each IFN. Fluorescence was analyzed 24h post-infection (hpi). In concordance with our results from Figure 1A, we found that only conspecific IFN was able to block viral replication in chicken and duck cells (Figure 1B), whereas all three avian IFNs were able to block viral replication in quail cells (Figure 1B). Mammalian universal IFN was only able to block viral replication in human A549 cells (Figure 1B). The data from the quail cells recapitulated results from earlier studies that found that chicken IFN could induce OASL activity in quail cells(26) and protect them from infection with vesicular stomatitis virus, *Salmonella typhimurium*, and avian bornavirus(26–28). Together, these data suggest that quail interferon receptors may be flexible receptors for IFNs from other species, whereas chicken and duck interferon receptors are more selective.

### Avian species exhibit species-specific transcriptomic responses to interferon and IAV infection

We sought to identify avian ISGs by looking for genes upregulated in chicken cells following conspecific IFN treatment. Here, we defined an ISG as a differentially expressed protein-coding gene with a fold change ≥2 relative to mock treated conditions and an adjusted p-value ≤0.05. Using these criteria, we identified 101 ISG in chickens following 4h of IFN treatment and 119 ISGs following 6h of IFN treatment (Figure 2A, Supplementary Figure 1A-B). The overlap between both timepoints was 43.8%, resulting in 153 unique ISGs. This number is consistent with previous studies identifying chicken ISGs that found between 62 and 332 ISGs(3, 29–31). To expand our inquiry to other avian species, we treated duck and quail cells with conspecific IFN to identify ISGs in those species as well. 204 ISGs were identified in duck and 81 in quail (Figure 2A, Supplementary Figure 1C-D).

**Figure 2.**
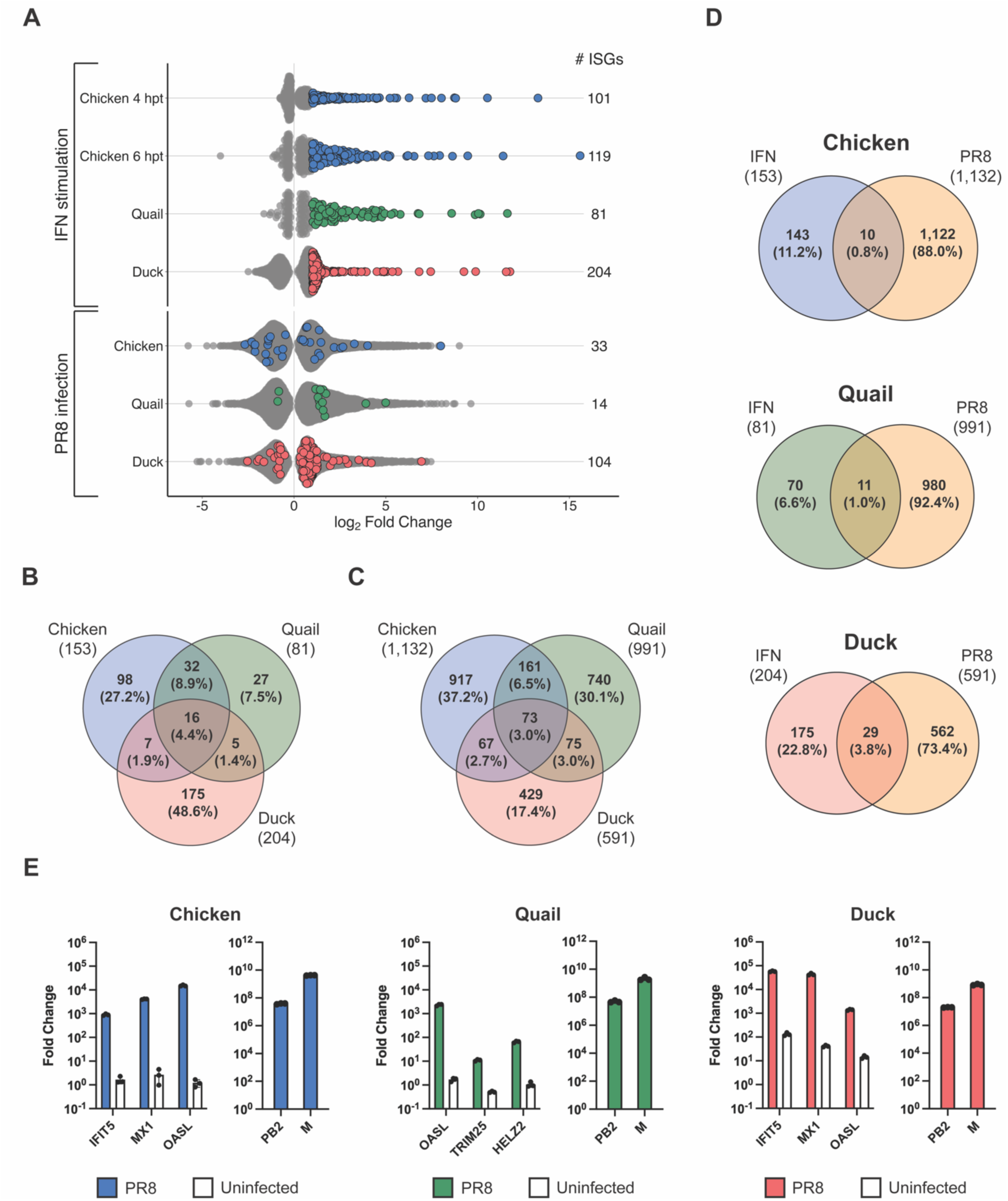
**A)** (top) Chicken, duck, and quail cells were treated with 200ng/mL of conspecific IFNα for either 4 (chicken and duck) or 6h (chicken and quail) before and transcriptomes profiled by RNAseq. Untreated cells were used as controls. ISGs were defined as protein-coding genes with a fold change ≥ 2 and an adjusted *p*-value < 0.05. Data are from a single independent experiment with five samples per group. **A)** (bottom) **C)** Chicken, duck, and quail cells were infected with PR8 at an MOI of 5. Cells were collected for RNA-sequencing 6h post infection. Genes with a fold change ≥ 2 and an adjusted *p*-value < 0.05 were considered upregulated. Red dots indicate ISGs identified in with conspecific IFN. Data are from a single independent experiment with five samples per group. **B)** Overlap between chicken, duck, and quail ISGs. Bold type indicates that that gene is also an ISG in humans **C)** Overlap between chicken, duck, and quail PR8-stimulated genes. **D)** Overlap between ISGs and PR8-stimulated genes for each species. **E)** Quantitative rtPCR for selected ISGs and the PB2 and M genes of PR8 in chicken, quail, and duck. PR8-infected samples from Figure 2C. N = 3 samples per group.

Having demonstrated that IFN activity is largely species-specific we next evaluated if the transcriptional response to IFN was also species-specific across the avian species tested. We compared the ISGs identified in each species and found little overlap between chicken, duck, and quail (Figure 2B), including only 16 ISGs shared between all three species. 48 ISGs were shared between chicken and quail, which are both galliform species and more closely related to each other than they are to ducks. We found fewer ISGs shared with the more divergent duck, with only 23 ISGs shared between chicken and duck, and 21 shared between quail and duck. Increasing the stringency of ISG calling to a fold change > 16 did not appreciably affect the number or percentage of ISGs that overlapped between species (Supplementary Figure 2A). Combining all chicken ISGs identified in our study, along with those identified in previous studies led to a total of 543 ISGs (Figure 3A). Comparing those chicken ISGs to a rationally curated list of 583 human ISGs(32, 33) resulted in an overlap of 6% (Supplementary Figure 2B). These results indicate thatmany ISGs in avian species may be species-specific.

**Figure 3.**
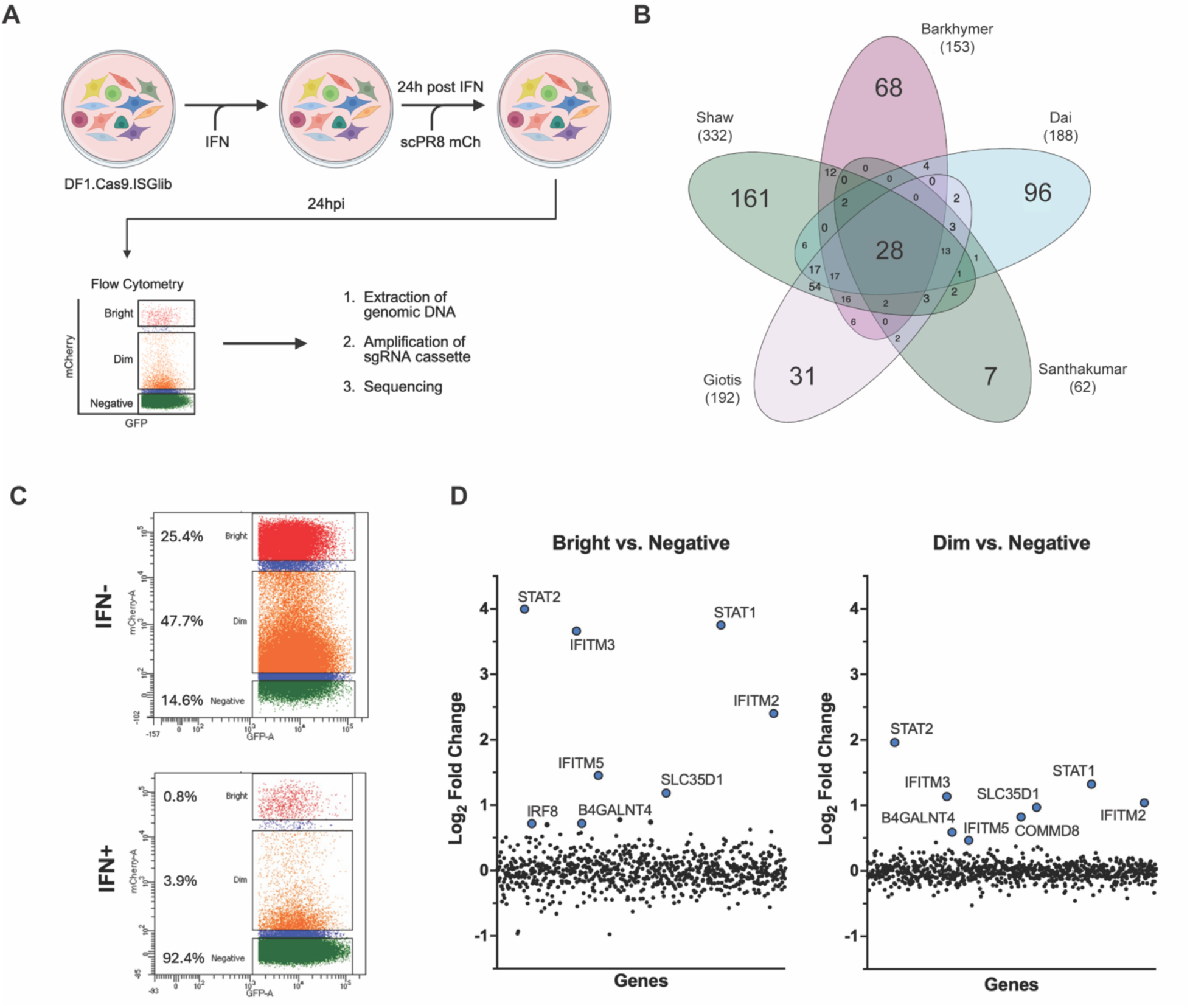
**A)** Schematic of CRISPR knockout screen. DF1.Cas9.ISGlib cells were treated with 200ng/uL chIFN 24h prior to being infected with scPR8-mCh at an MOI of 5. 24h after infection cells were sorted on fluorescence and processed for amplicon sequencing. **B)** Overlap of chicken ISGs identified in five separate studies. Diagram made using InteractiVenn(73). **C)** Screened cells from (D) were sorted by flow cytometry to isolate iRFP+ GFP+ cells, which were then sorted based on mCherry fluorescence into mCh “Bright”, mCh “Dim”, and mCh “Negative” populations. **D)** Top hits from screen. Three independent screens were pooled for analysis. Log2 fold-change of each ISG is shown. Genes are arranged randomly along the x-axis. Colored and labeled genes are statistically significant (Adjusted P-value< 0.05).

We also evaluated transcriptional responses to IAV infection in birds by infecting chicken, quail, and duck cells. Cells from these species were infected with PR8 and upregulated genes evaluated 6h later. We found that chicken and quail had 1,132 and 991 virus upregulated genes, respectively, whereas only 591 genes were upregulated 6hpi in duck (Figure 2A, Supplementary Figure 3A-C). As with the ISGs, we found very little overlap between the PR8-stimulated genes of each species (Figure 2C). To determine whether infection with PR8 also led to the upregulation of ISGs, we compared the total number of ISGs identified earlier with the PR8-stimulated genes of each species (Figure 2D). We found that chicken and quail had an overlap of 0.8% and 1.0% between the two gene sets, respectively, indicating that ISGs were not generally upregulated early during PR8 infection. This is supported by qRT-PCR data showing that selected ISGs were not upregulated in the samples used to generate the lists of PR8-stimulated genes (Figure 2E; left). In duck, however, we found a 3.8% overlap between ISGs and PR8-stimulated genes, almost four times higher than the overlap in chicken and quail. This is also supported by quantitative PCR data showing that selected ISGs were upregulated in duck samples, though the difference in fold change of the ISGs between the IFN-treated and the PR8-infected samples was about 3 logs. To ensure that all our samples were infected with PR8 to an equivalent degree, a quantitative PCR was performed targeting viral mRNA. We observed high levels of viral mRNA in all samples (Figure 2E; right), indicating that a difference in level of infection was not the cause of differential ISGs upregulation across species.

### CRISPR screen identifies several genes with potential antiviral activity against PR8

To determine which ISGs are capable of restricting IAV in birds, we performed a pooled CRISPR knockout screen of chicken ISGs (Figure 3A). To compile the list of ISGs to use for the screen, we combined chicken ISGs identified at both 4 and 6hpt with chicken ISGs identified in previous studies(3, 29–31) and generated a list of 543 chicken ISGs (Figure 3B). Using this list, an sgRNA library was generated consisting of five sgRNAs per gene along with 275 nontargeting control sgRNAs. The sgRNAs were cloned into a retrovirus-based plasmid expressing an infrared RFP fluorescent marker (miRFP670) and a blasticidin resistance gene. The sgRNA-encoding retroviruses were then produced by transfection into HEK293T cells and titered on chicken DF-1 cells. To knock out genes of interest in DF-1 cells, a DF-1 cell line was used that constitutively expresses Cas9 fused to GFP (DF1.Cas9)(5). The knockout cell library was made by transducing the DF1.Cas9 cells with the sgRNA-encoding retrovirus pool at an MOI of 0.5, to reduce the likelihood of double integrants. This was done three separate times to create independent libraries of cells. To perform the screen, we treated cells with 200ng/mL chicken IFN 24h prior to infection with scPR8-mCh at an MOI of 5. This resulted in a ten-fold reduction of replication (Figure 3C). Cells double positive for iRFP and GFP were isolated by flow cytometry and sorted into bright, dim, and mCherry-negative populations at 24hpi (Figure 3C). Amplicon sequencing to identify enriched sgRNAs was performed following extraction of genomic DNA and PCR amplification of the sgRNA cassette.

We performed a Robust Rank Aggregation analysis using MAGeCK and identified nine hits in this initial screen (Figure 3D). We analyzed both the ‘Bright’ and the ‘Dim’ group, using the ‘Negative’ group as the comparison control. STAT1 and STAT2 are known to be involved in the induction of ISGs in mammals, whereas mammalian IRF8 is important for pro-inflammatory signaling in dendritic cells(34). In mammals, STAT1 and STAT2 associate with IRF9 to induce the expression of ISGs(35). Chickens lack an IRF9 gene, making it unclear which transcription co-factor enables ISG expression(36). The identification of IRF8 as a hit in our screen along with STAT1 and STAT2 is the first evidence that IRF8 may compensate for the lack of IRF9 in chickens in JAK/STAT signaling. The other hits identified were B4GALNT4, COMMD8, IFITM2, IFITM3, IFITM5, and SLC35D1. IFITM2 and IFITM3 have previously been shown to restrict IAV in chicken cells(14), and IFITM3 has been purported to be responsible for 40%-70% of the restrictive activity of ISGs in human cells(37). IFITM3 functions as an entry inhibitor for IAV. It localizes to late endosomal membranes and prevents IAV RNPs from entering the cytoplasm by blocking the formation of the fusion pore(38, 39). IFITM2 has also been shown to restrict IAV in both humans and chickens, though not as effectively as IFITM3(14). In humans, IFITM5 is only expressed in bone tissue and is not thought to have any immunity related function(40). Chicken IFITM5 has been previously shown to restrict retroviruses pseudotyped with glyocoproteins from filoviruses (e.g. Ebolavirus and Marburg virus) but not influenza viruses(41). The roles of B4GALNT4, COMMD8, and SLC35D1 in IAV restriction have not been investigated.

### IFITM3 is the primary antiviral factor against PR8 in chicken cells

To validate the hits from our screen, we generated three independent knockout chicken DF-1 cells targeting these genes and a non-targeting control specific for *E. coli* LacZ. We included several genes related to our hits including IFITM1, IFITM10, and ATHL1. IFITM1 and IFITM10 were not included in our CRISPR screen library because they were not identified as ISGs by RNA-seq but given genetic and structural similarities to other IFITM family members, we decided to include them in our validation experiments. In addition, we noted that in addition to IFITM2, IFITM3, and IFITM5 identified as hits, B4GALNT4 was enriched in our screen. B4GALNT4 is located telomeric and adjacent to the IFITM gene family on chicken chromosome five (Figure 4B). Out of curiosity regarding antiviral function of genes neighboring the IFITM locus, we included ATHL1 that is located centromeric and adjacent to the IFITM gene family. ATHL1 (also known as PGGHG) is not an ISG in chickens or humans. MRGBP was also included in our validation experiments because it was enriched in the ‘Dim’ group though it was not enriched in the ‘Bright’ group.

**Figure 4.**
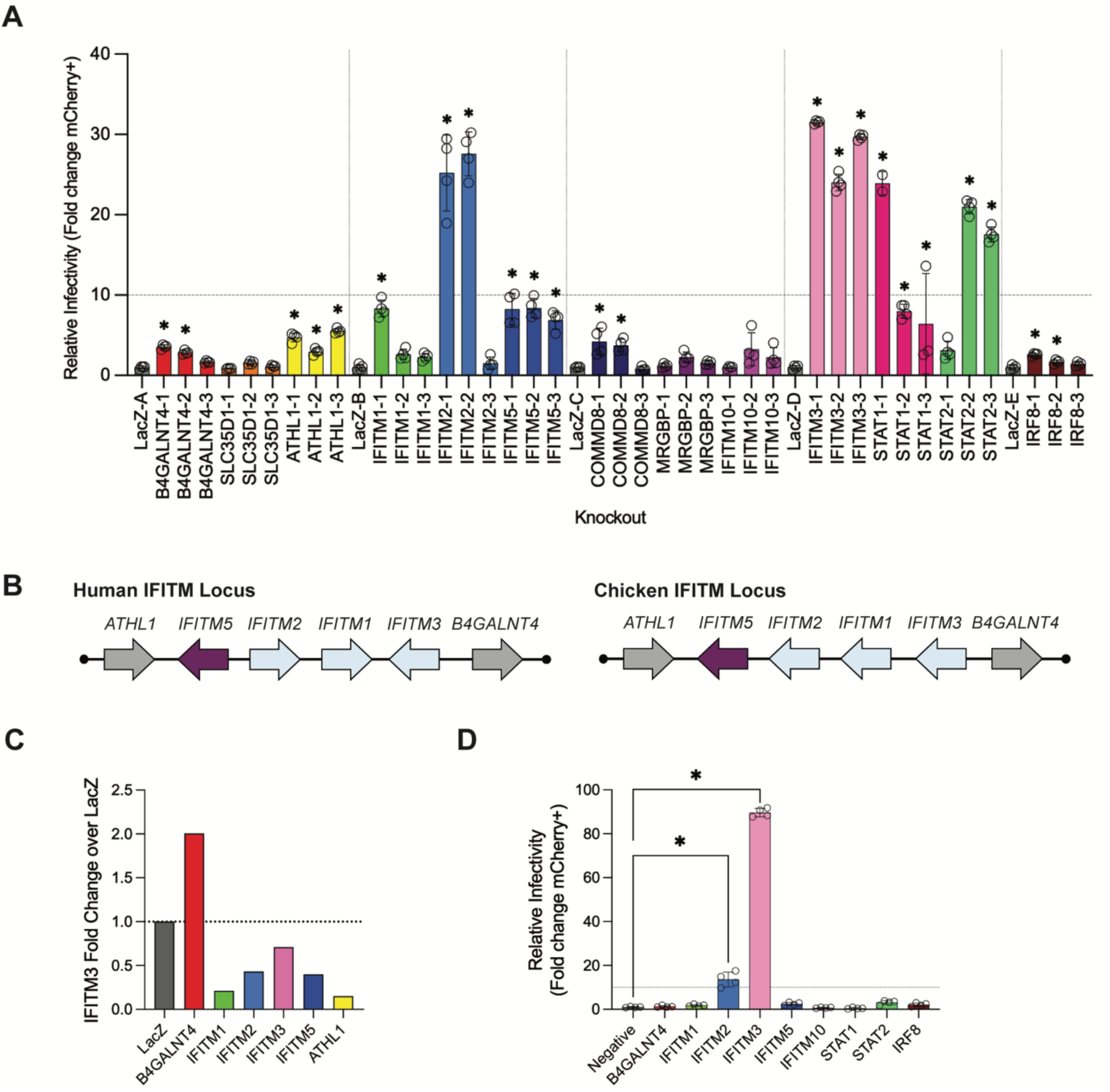
IFITM3 is the primary restrictor of PR8. **A)** Polyclonal knockout DF-1 cells were treated with 200ng/mL chicken IFNα for 24 hours following infection with scPR8-mCh at an MOI of 5. Cells were analyzed for mCherry fluorescence 24 post infection. Percent infection in LacZ knockout cells was normalized to 1, with the values for the other genes given as fold change relative to that. The experiment was done in four separate rounds (separated by dotted vertical lines), with each round having its own LacZ control. N = 4 samples per group. **B)** Schematic of the human and chicken IFITM locus. **C)** qPCR for IFITM3 expression performed on LacZ, B4GALNT4, IFITM1, IFITM2, IFITM3, IFITM5, and ATHL1 knockout cells following IFN treatment. One biological replicate and four technical replicates per group. Data show fold change of IFITM3 expression over IFITM3 expression in LacZ knockout cells. **D)** DF-1 cells were transfected with siRNA for indicated genes 24h prior to treatment with 200ng/mL chicken IFNα. Cells were infected with scPR8-mCh at an MOI of 5 at 24h after IFN treatment and mCherry fluorescence was measured 24h following infection. N = 4 samples per group. Statistical analyses for **(A)** and **(D)** done using one-way ANOVA with Dunnet’s multiple comparisons. * indicates p-value < 0.01.

We treated knockout DF-1 cells with chicken IFN for 24h followed by infection with scPR8.mCh for 24h (Figure 4A) prior to fixation, imaging, and quantification of percent infection. Three sgRNAs were used per gene. These infections largely validated our hits using the same fluorescent virus used in the original screen. The CRISPR knockout validation identified IFITM2, IFITM3, STAT1, and STAT2 as strong restrictors of IAV infection, with an over 20-fold increase in cells infected. Less extreme but still statistically significant were B4GALNT4, ATHL1, IFITM1, IFITM5, and COMMD8.

Knockout of ATHL1 modestly increasing scPR8-mCh infectivity raised questions that the genetic manipulation of the IFITM genes might affect other genes in the locus, making it difficult to discern which genes were actually restricting IAV. To determine if this was the case, *IFITM3* mRNA expression was measured in IFITM1, IFITM2, IFITM3, IFITM5, B4GALNT4, and ATHL1 knockout cells following IFN stimulation. The expression of IFITM3 in these cells was compared to that in control LacZ knockout cells, which should express wild-type levels of these genes. In IFITM1, IFITM2, IFITM3, IFITM5, and ATHL1 knockout cells, *IFITM3* mRNA expression levels were lower than in the LacZ control cells, indicating that the knockout of these genes might impact *IFITM3* expression (Figure 4C). IFITM3 expression levels in B4GALNT4 knockout cells was double that in control cells, indicating that B4GALNT4 knockout does not negatively impact IFITM3 expression.

To overcome this caveat we used an independent approach and employed siRNA knockdown, which targets the mRNA of genes and is therefore unaffected by genomic proximity. Here, IFITM3 was again the strongest gene that restricted IAV when the experiment was done using scPR8.mCh (Figure 4D). These results indicate that there may be some form of genetic or chromosomal modulation occurring during knockouts within the IFITM locus potentially resulting in false positives. However, we also observed that while STAT1 and STAT2 strongly restricted IAV in our knockout validation, STAT1 and STAT2 knockdown had limited effects on IAV replication. This is likely due to lack of complete depletion of STAT1 protein and the need for a minimal amount of protein to induce a restrictive effect. This points to caution in drawing conclusions based on siRNA results alone and highlights the importance of using orthogonal methods of validation.

### Avian IAV is restricted by overlapping and unique chicken ISGs

PR8 is a laboratory strain that has been serially passaged and adapted to both mouse lungs and chicken eggs. Therefore, we repeated our flow-based screen using the avian virus A/guinea fowl/Hong Kong/WF10/99(H9N2) (WF10). We found that chicken IFNα was less effective at restricting WF10 than PR8 (Figure 5A). Whereas 24h of IFN treatment prior to PR8 infection resulted in a 10-fold reduction in IAV replication, WF10 replication was only inhibited 30% or less when measured as a binary of infected versus uninfected. However, when the percentage of cells in the “Bright” group were compared with and without IFN treatment, a much greater level of restriction is observed (Figure 5A). This indicates that for WF10, while chicken IFNα is potentially less effective at preventing the virus from entering the cell and establishing an infection, it can restrict the transcription of viral mRNA within the cell. The difference in IFN-mediated restriction between WF10 and PR8 might be due to PR8 is grown in chicken eggs prior to IFN signaling turning in in chicken embryos, leading to a lack of selective pressure on the virus to overcome IFN restriction.

**Figure 5.**
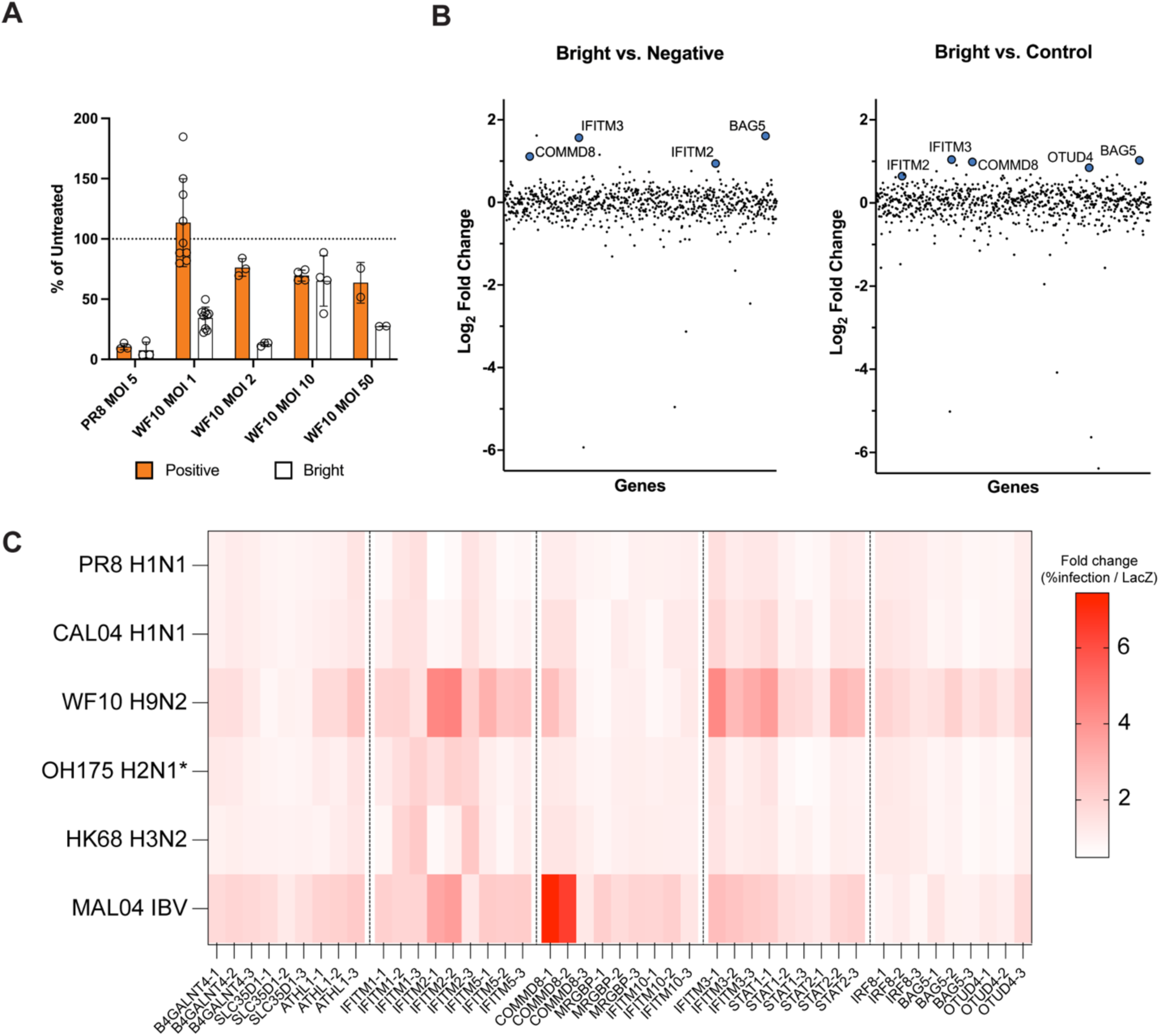
WF10 is inhibited by a similar set of genes as PR8. **A)** Percent of infected or ‘Bright’ cells based on flow cytometry. **B)** Top hits from screen. Three independent screens pooled. Log2 fold-change of each ISG is shown. Genes are arranged randomly along the x-axis. Colored and labeled genes indicate adjusted p-value < 0.05. **C)** Polyclonal knockout DF-1 cells were treated with 200ng/mL chicken IFNα for 24 hours following infection with the indicated viruses at an MOI of 5. Cells were analyzed for mCherry or NP+ fluorescence at 24 post infection. Percent infection in LacZ knockout cells was normalized to 1 (white), with the values for the other genes given as the fold change relative to that (red scale). The experiment was done in five separate rounds (separated by dotted vertical lines), with each round having its own LacZ control. N = 4 samples per group.

The results of our flow-based screen using WF10 also identified IFITM1, IFITM2, and COMMD8 as hits and additionally identified the genes BAG5 and OTUD4 (Figure 5B). Bright and Dim groups were compared to both Negative and Control (uninfected and untreated) groups. BAG5 is a co-chaperone protein of heat shock protein 70(42). OTUD4 is a deubiquitinase implicated in MyD88/NFκB signaling(43) as well as promotion of MAVs signaling(44). We validated these hits as above using scPR8-mCh (Figure 4A). To determine how broadly all the putative hits restrict IAVs, we infected individual knockout DF-1 cells with wild-type PR8, three mammalian origin viruses, and two avian origin viruses (Figure 5C). wild-type PR8 yieled similar results as our scPR8.mCh infections in Figure 4A, albeit with somewhat lower fold changes relative to LacZ control DF-1 cells. Interestingly, COMMD8 was identified as a strong restrictor of influenza B virus. We also found that OTUD4 and BAG5 preferentially restricted avian influenza viruses. Together the individual KO cells validated the results from the screen.

## DISCUSSION

In this study, we sought to identify ISGs that restrict IAV infection in avian cells. We first evaluated the antiviral landscape in chicken, quail, and duck fibroblasts, and in general, found substantially fewer than has been established for humans (∼500-1000)(32, 33). However, combining chicken ISGs from various studies revealed 543 chicken ISGs. There were differences in cell type and time point used across studies, both of which can impact ISGs detected. Studies comparing DF-1 cells to chicken embryonic fibroblasts have found that DF-1 cells have constitutively elevated levels of SOCS1, a protein that dampens the cell’s proinflammatory responses(45). This could feasibly result in fewer ISGs being identified in experiments using DF-1 cells. It is becoming increasingly more appreciated that there are cell and tissue specific ISGs(46). Additionally, species-specific expression or function of ISGs that control influenza (KHNYN/ZAP, MX1, BTN3A3) can have impacts for cross species replication and may control emergence of viruses into new species(47). To this point, little overlap was observed in the transcriptomic responses to IFN and IAV infection between chicken, quail, and duck cells. This is consistent with a study evaluating ISGs in ten species, including chickens, that found only 62 common ISGs despite most species having hundreds of genes upregulated(3). In addition to the presence and absence of ISGs, there can be substantial variation in genomic sequence across species and ISGs are under heavy positive selection when compared to other genes(3, 4). Chickens and ducks diverged 83 million years ago, and chickens and quails 33 million years ago, leaving sufficient time for positive selection to generate divergent ISG-omes. Indeed, chickens and quails have more ISGs in common than chickens and ducks, likely due to these animals having diverged more recently. In addition to genetic differences in antiviral genes mammals and avians can have substantially different physiologies. The basal temperature of most birds is higher than mammals and temperature can impact influenza replication and induction of immune responses (41481744, 41308154). Further antiviral genes may have been selected for function at higher fever temperature (41379117). The screens and validations in this study were performed at 37C and it would be interesting in future studies to determine if there are similar and/or different hits at 39 or 41C.

The main goal of this research was to determine which chicken ISGs control IAV infection. We discovered that IFITM3 is the primary ISG involved in restricting IAV in chicken cells capable of at least 30-fold suppression of IAV replication in a single infection cycle. At first, this may seem surprising, as the chicken ISG-ome potentially contains hundreds of genes, however, recent data suggests that a small subset of ISGs may be responsible for the majority of IFN’s restrictive function against any given virus(48). Accordingly, it is feasible that only a few ISGs are responsible for IAV restriction in chickens. Most other ISGs that restrict IAV in humans, such as PKR, TRIM25, MOV10, and GPB, have unknown restrictive functions in birds and our data suggest that they do not play a major role in restricting IAV in chicken cells. In such cases, some ISGs may operate but are masked by the dominant inhibitory effects of major restriction factors. IFITM3 has already been shown in a previous siRNA screen to be responsible for the majority of the IFN-mediated IAV restriction(37). More recently, a study revealed that human and mouse IFITM3 can restrict a wide range of human, swine, and avian IAV viruses, and that IFITM3 deficiency lowers the minimum infectious dose threshold in mice(49). IFITM3 deficiencies in humans have also been linked to increased susceptibility to IAV infection and worse outcomes(50–52).

Surprisingly we also found hits that were avian strain specific, BAG5 and OTUD4. OTUD4 is a deubiquitinase that interacts with MAVS and other proteins to promotes antiviral signaling during viral infection(43, 44), in our screen this may help to increase antiviral gene expression or OTUD4 may directly interact with viral proteins(53). BAG5 is a co-chaperone to heat shock protein 70 (HSP70). The role of HSP70 in IAV infection is unclear. HSP70 may interact with IAV RNPs and overexpression of HSP70 in IAV-infected A549 cells may decrease viral titer as well as viral RNA transcription and protein expression(54). In addition, overexpression of HSP70 can inhibit the nuclear export of viral proteins(55). However, IAV polymerase activity might also be decreased in cells with augmented HSP70 expression(56), indicating that the role of HSP70 in IAV infection is incompletely characterized. We speculate that BAG5 and OTUD4 may exhibit more pronounced restriction against WF10 allowing their enrichment in our screen, due to strain-specific sensitivity to WF10. Alternatively, we posit that flow-based enrichment of WF10 (via surface-exposed HA epitope tagged HA protein expressed in infected cells by antibody staining based flow cytometry) rather than intracellular mCherry expression generated from the HA vRNA segment could augment hit enrichment.

Together, this study demonstrates that IFITM3 is the ISG most responsible for restricting IAV in chicken cells. We also uncovered additional genes with varying abilities of restricting IAV. Further exploration of hits as well as other avian ISGs should be a priority in the future, as they could provide important information about the control of IAV in birds that may mitigate the impacts of avian influenza and help to uncover biology of viruses at the cross-species interface.

## METHODS

### Cell culture

A549 (male human lung carcinoma, ATCC, CCL-185), HEK293T (male human epithelial kidney, ATCC, CRL-3216), DF-1 (chicken embryonic fibroblast, ATCC, CRL-12203) cells, and their derivatives were cultured in Dulbecco’s Modified Eagles Medium (DMEM, Gibco) supplemented with 10% Fetal Bovine Serum (FBS, ThermoFisher) and 1% penicillin/streptomycin solution (ThermoFisher). CCL-141 cells (duck embryonic fibroblast, ATCC) were cultured in Eagle’s Minimum Essential Medium (EMEM, ATCC) containing 10% FBS and 1% penicillin/streptomycin solution. QT-6 cells (quail fibroblast-like ATCC, CRL-1708) were cultured in Ham’s F-12K Medium (ThermoFisher) containing 10% FBS, 1% penicillin/streptomycin solution, and 10% tryptose phosphate broth (ThermoFisher). All cell lines were obtained from ATCC, maintained at 37°C with 5% CO_2_, and used without extensive passaging.

### CRISPR/Cas9 knockout library generation

An sgRNA library of 543 chicken ISGs (2715 total on-target sgRNAs; ∼5 sgRNAs/gene) and 275 nontargeting control sgRNAs was designed by GenScript Biotech in collaboration with the authors using the Synthego CRISPR sgRNA design tool or RGEN Cas-Designer(57). sgRNA library sequences are provided in Supplemental Table 1. CRISPR library retrovirus pools were generated as described above and titered by serial dilution on DF-1.Cas9 cells measuring miRFP670 fluorescence using a Cytation 5 Cell Imaging Multimode Reader (Agilent, Biotek). To generate an ISG knockout library in DF-1 cells, six million DF-1.Cas9 cells were transduced at an MOI=0.5 infectious units per cell to ensure one sgRNA transduction target per cell and subsequently selected with up to 8µg/mL blasticidin starting at 48 hours post-infection and maintained throughout continuous cell culture. We generated three independent ISG knockout cell libraries in DF-1.Cas9 cells that were expanded and cryo-preserved for future use. For screening ISGs, at least three million cells (Cas9+sgRNA+) were plated in 10-cm plates and treated with 200ng/mL of recombinant chicken IFN-alpha (Bio-Rad PAP004) after 24 hours. Following 24 hours of chicken IFN administration, cells were washed with PBS, infected with IAV at MOI=5 in infection media (1x PBS, 2.5% bovine serum albumin, 1% calcium/magnesium) for one hour, washed with PBS, and incubated in media overnight until processing for fluorescence activate cell sorting.

### Plasmids

CRISPR/Cas9 editing of avian cells was performed by sequential retroviral transduction of Cas9-GFP expression vector followed by single guide RNA (sgRNA) expression vector. Briefly, guide RNAs were designed from the literature, Synthego design tool, or RGEN Cas-Designer, complementary oligos with Esp3I restriction enzyme compatible ends were annealed, and subsequently ligated into pMLV.miRFP670.iU6sgRNA. Retroviral Cas9-GFP expression vector and pMLV.mIRFP670.iU6sgRNA have been described previously(5). Plasmid library encoding 2715 sgRNAs for 543 ISGs (∼5 sgRNAs per gene) and 275 non-targeting / PAM-less control guides was designed, cloned, expanded, and sequenced by GenScript Biotech with iterative oversight from the authors. Other DNA constructs were sequence confirmed by whole plasmid sequencing performed by Plasmidsaurus using Oxford Nanopore Technology or Sanger sequencing performed by Eurofins Scientific or GeneWiz from Azenta. sgRNA sequences used for individual knockouts are provided in Supplemental Table 2.

### Retroviral transductions and stable cell generation for CRISPR/Cas9 knockout

Simple retroviruses for transduction were generated by transfecting HEK293T cells in 100mm dishes with 4.5µg vector plasmid(58, 59), 4.5µg pMD-gagpol(60), and 1µg pMD-VSVG(61) using Transit-LT1 (Mirus Bio), changing media after 24 hours, and harvesting culture supernatant at 48 hours post-transfection followed by 0.45µm syringe filtration. Transducing virus preps were aliquoted and stored at -20°C. Stable cells were generated as previously described(62). DF-1 cells were transduced with simple retrovirus Cas9-GFP (puromycin-resistant) to generate DF-1.Cas9 cells. DF-1 cell knockouts were maintained as pools stably expressing Cas9 and sgRNAs (miRFP670+ and blasticidin-resistant). For single gene knockout cells, approximately 2500 target cells were seeded into a 96-well flat bottom plate, allowed to adhere overnight, and 20-200µL of transducing viral supernatant with up to 10µg/mL polybrene added to each well. After two days in culture, transduced cells are washed with PBS, and fresh media containing antibiotic (2µg/mL puromycin, 2-8µg/mL blasticidin; GoldBio, Inc) added. Surviving cells were grown and expanded in the ongoing presence of antibiotic.

### Influenza viruses

Single-cycle PR8 mCherry (scPR8.mCh) viruses have been described previously(25). Briefly, scPR8.mCh were rescued by plasmid-based transfection using Lipofectamine 3000 into HEK293T cells (500ng of seven bicistronic PR8 plasmids [PB2, PB1, PA, NP, NA, M, NS], 500ng pCAGGS-HA, and 500ng pPol.HA-Pack.mCherry) and subsequently amplified in MDCK cells stably expressing PR8 HA. Wild-type PR8, CAL04 (A/California/04/2009(H1N1))(63), HK68 (A/Hong Kong/01/1968(H3N2)), OH175 using PR8 HA and NA (A/Green-winged teal/Ohio/175/1986(H2N1))(64), WF10 encoding an HA epitope-tagged HA (A/guinea fowl/Hong Kong/WF10/1999(H9N2))(65), and MAL04 (B/Malaysia/2506/2004)(66) have been described. Virus titers were determined by plaque assay in MDCK cells as previously described(67) or by flow cytometry in the case of WF10.

### Influenza single-cycle infections

For single-cycle influenza virus infections, cells were plated and allowed to grow overnight for up to 24 hours. Cells were treated with IFN (200ng/mL for chicken, duck, and quail IFN or 1000 units/mL for universal IFN) by replacing media and returned to incubation at 37°C for indicated time points. Subsequently, cells were infected at an MOI of 5 infecting a confluent culture of cells with at least three independently infected well replicates. For infection, cells were washed with PBS prior to addition of infection media (1x PBS, 2.5% bovine serum albumin, 1% calcium/magnesium). Viral stocks were diluted to appropriate MOI in infection media, transferred to wells, and cells were incubated for 1 hour at 37°C. Then, cells were washed once with PBS prior to addition of cell culture media. For CRISPR screen and hit validation infections, TPCK-resistant trypsin was excluded from infected cultures to prevent spreading replication.

### siRNA knockdowns

For transient knockdown by siRNA, cells were plated and transfected with mock, 100nM control, or 100nM pooled duplexed siRNAs targeting Gallus gallus mRNA transcripts using the HiPerFect transfection kit according to manufacturer’s instructions (QIAGEN). Negative control DsiRNA (IDT Cat# 51-01-14-03), DsiRNA targeting *Renilla* luciferase (IDT Cat# 51-01-08-22), and mock transfected wells were used for comparison. siRNAs were designed using IDT dsiRNA design tool and sequences are provided in Supplemental Table 2.

### Quantitative Reverse Transcription PCR

First strand cDNA synthesis was done using 1ug of RNA using the SuperScript IV First Strand Synthesis system with Oligo d(T) primers (Thermofisher). RT-qPCR was performed using iTaq Universal SYBR Green Supermix (Bio-Rad) using a CFX96 Real-Time System C1000 Touch Thermocycler (Bio-Rad). Expression levels were quantified by the delta-delta Ct method relative to a loading control gene (TUBA4A for human, chicken, and duck, β-actin for quail). Primer sequences are listed in Supplemental Table 4.

### RNA Sequencing

RNA sequencing was performed using the NovaSeq 6000 or the NovaSeq X Plus with 150 paired-end reads. Library prep was done using the Kapa Hyper Stranded mRNA library kit (Roche). Sequencing results were trimmed using Trimmomatic version 0.39(68). Trimmed reads were aligned to the genome using Star version 2.7.8a(69). Reads were counted using HTseq version 1.99.2(70). The GRCg7b chicken genome, the CAU_duck1.0 duck genome, and the Coturnix_japonica_2.0 quail genome were used for alignment. Differential gene analysis was performed in R using DESeq2(71) comparing IFN-treated or PR8-infected conditions to the untreated condition. Differentially expressed genes were defined as protein-coding genes with a Log_2_ fold change ≥ 1 (representing at least 2-fold increase in expression) and an adjusted p-value <0.05. For IFN-treated samples, ensembl IDs with no associated gene name were identified using BLAST. This was not done for PR8-infected samples due to the high number of unnamed genes.

### Fluorescence microscopy

Fixed-cell images were obtained on a BioTek Cytation 5 (Agilent) using a 4X (NA=0.13, #1220519), 20X (NA=0.40, #1220517), or 40X (NA=0.6, #1220544) Plan Fluorite objective and LED-filter cube sets for DAPI (excitation = 377±50nm; emission = 447±60nm; #1225007 and #1225100), GFP for mNeonGreen (excitation = 469±35; emission = 525±39; #1225001 and #1225101), TRITC for mCherry and AlexaFluor568 (excitation = 556±29; emission = 600±37; #1225012 and #1225125), and Cy5 for miRFP670 (excitation = 620±40; emission = 685±40; #1225105 and #1225005). We quantified single-cycle IAV infections by imaging and used the BioTek Gen5 software to calculate percent infectivity using scPR8 (MOI=5) encoded mCherry as a readout for infectivity (# TRITC+ cells / # DAPI-stained nuclei). To quantify wild-type / non-mCherry non-replicating infections, we performed immunofluorescence on infected cells (MOI=5 without TPCK trypsin to block virus transmission) fixed with 4% paraformaldehyde at 24 hours post-infection. Briefly, we subjected cells to permeabilization (PBS with 0.25% Triton X-100) for 10 minutes, blocking (PBS with 3% bovine serum albumin), primary antibody staining with mouse anti-NP (BEI, NR-43899, 1:500) in blocking buffer (3% bovine serum albumin) for one hour, secondary antibody staining with goat anti-rabbit AlexaFlour-568 (Invitrogen, #A-11031, 1:500), and nuclear counterstaining with DAPI (Sigma, D9542, 100ng/mL). Titering of chicken ISG sgRNA MLV library was done via serial dilution of the virus on DF-1 cells in a 96-well plate. Percent of cells infected was determined by miRFP670 fluorescence, which was then used to calculate the titer.

### Fluorescence activated cell sorting

For samples infected with scPR8-mCh, sorting was done after cells were washed and resuspended in PBS + 1% BSA prior to sorting. Sorting was done on a Cytek Aurora CS machine. For samples infected with WF10 (MOI 2), cells were washed with PBS, blocked in 5% milk in PBS, and stained using an anti-HA TRITC-conjugated antibody (Millipore-Sigma H9037, 1:140). gDNA was extracted from each sorted population as well as from unsorted, untreated, and uninfected controls using DNeasy Blood and Tissue DNA purification kit (QIAGEN). PCRs were then performed on the entire 100uL eluant from each sample using four 50uL volume PCR reaction mixes using Phusion polymerase (NEB). Primers were designed to amplify the sgRNA cassette and generate a 214bp amplicon (Forward: 5’ TTCCATGGACGCTACCGGTC, Reverse: 5’ GGACTATCATATGCTTACCG). The PCR reactions were run on an agarose gel and bands of the correct size were cut out. The DNA was extracted from the gel using the Gel and PCR Clean-up kit (Macherey-Nagel).

### DNA amplicon sequencing and analysis in ISG screens

Amplicons were then sequenced using the MiSeq i100 with at least one million reads per sample. Library prep was done using the Kapa Hyper Prep kit (Roche). Enriched genes were identified using MAGeCK(72). Hits were defined as genes enriched in the treatment group compared to the control group with an adjusted p-value ≤ 0.05.

## Statistical analyses

Statistical analyses were done using a one-way ANOVA with Dunnet’s multiple comparisons in GraphPad Prism version 10.

## Data availability

Sequence data were deposited and are available as FASTQ files in the NCBI sequence read archive under BioProject no PRJNA1461921. RNAseq count data are available under GEO accession numbers GSE330214.

## ACKNOWLEDGEMENTS

We wish to thank members of the Langlois labs for helpful conversations. We wish to acknowledge and thank Andrew Mehle for the A/green-wing teal/Ohio/1751986(H2N1) OH175/S009 plasmids, Peter Palese for HK68, Nicholas Heaton for MAL04, and Anice Lowen for tagged WF10. This work was supported by NIAID Centers for Excellence in Influenza Research and Response (CEIRR - https://www.ceirr-network.org) grant NHH75N93021C00017 and R01 AI148669 to RAL, F32-AI147813 to JTB, T32 AI007313 to AJB, and T32 HL007741 to FKS. We thank the University of Minnesota Flow Cytometry Resource Facility and the DNA Services team at the Roy J. Carver Biotechnology Center, University of Illinois at Urbana-Champaign for technical support for RNA and amplicon sequencing.

---

**Supplementary Figure 1.**
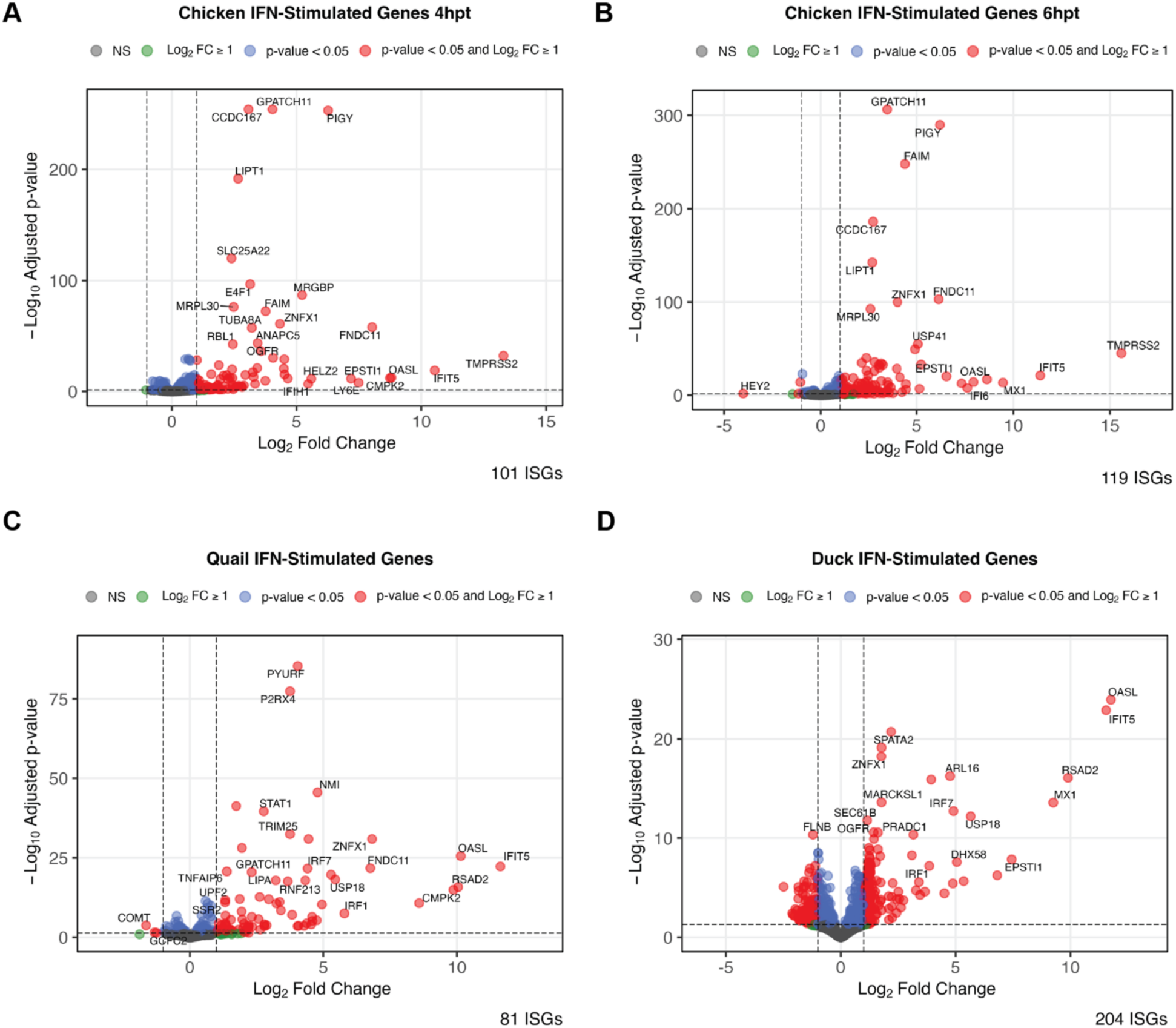
Interferon-stimulated genes are upregulated in response to conspecific IFNα treatment. Chicken cells (DF-1) were treated with 200ng/mL of recombinant chicken IFNa for 4h **(A)** or 6h **(B)** before being collected for RNA sequencing. **C)** Quail cells (QT6) were treated with 200ng/mL of quail IFNa for 6h before being collected for RNA sequencing. **D)** Duck (CCL-141) cells were treated with 200ng/mL of duck IFNa for 4h before being collected for RNA sequencing. N = 5 samples per group.

**Supplementary Figure 2.**
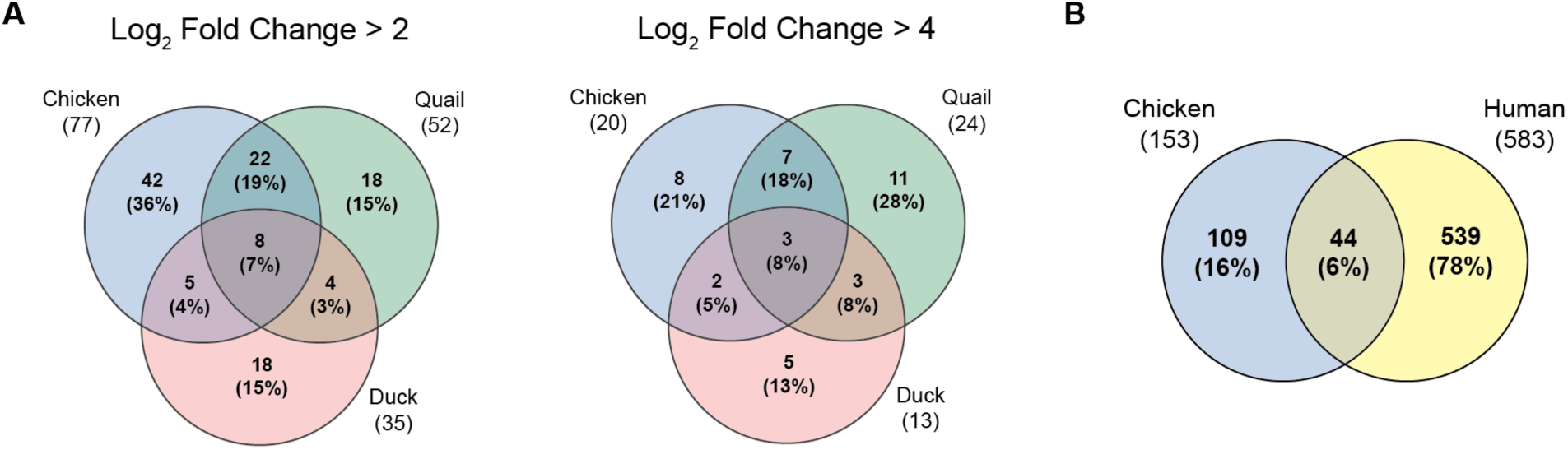
**A)** Overlap between chicken, duck, and quail ISGs, when the cutoff is Log2 fold change > 2 (foldchange > 4) and Log2 fold change > 4 (fold change > 16). **B)** Overlap between chicken ISGs identified in this study and human ISGs.

**Supplementary Figure 3.**
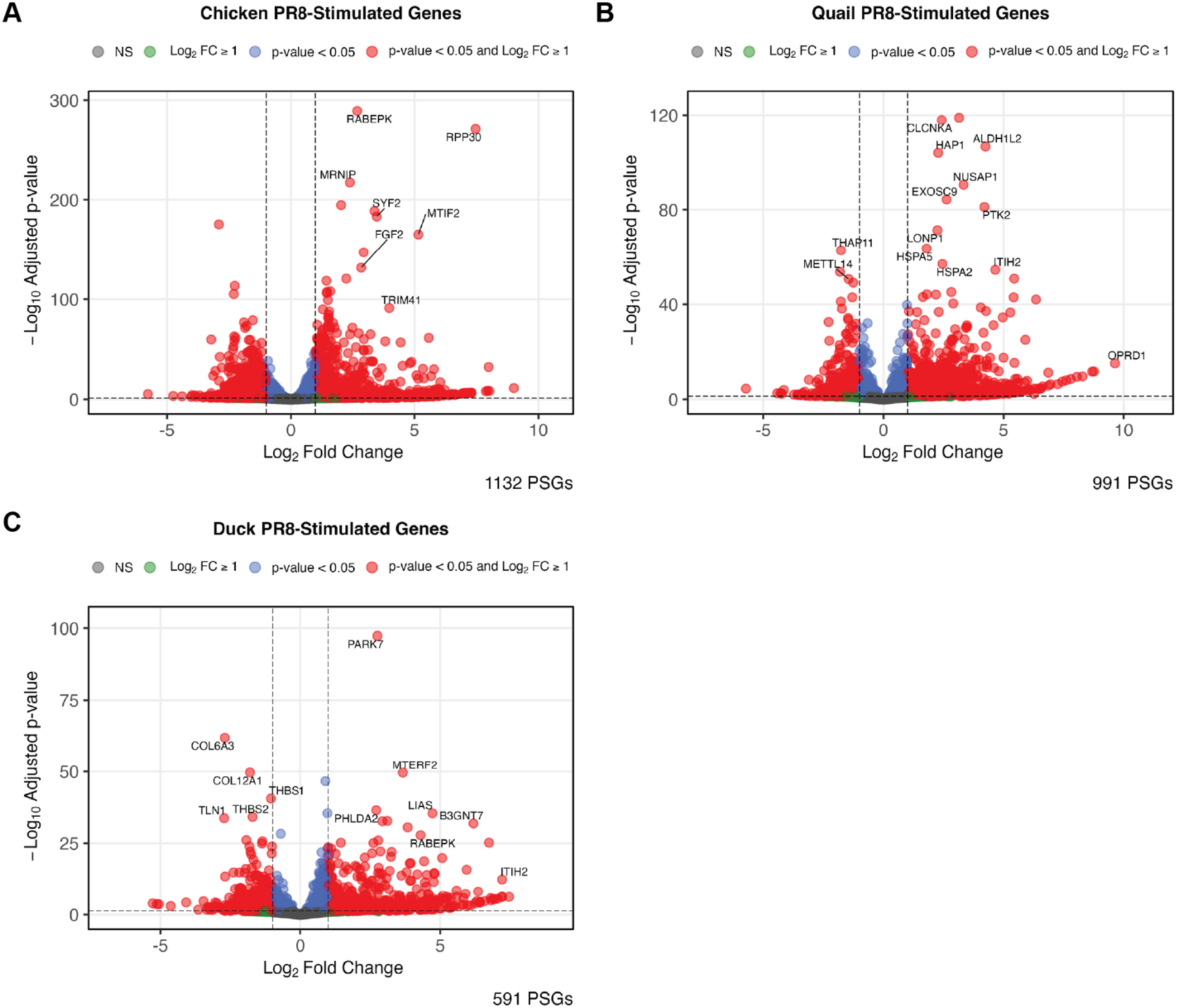
Genes upregulated during PR8 infection in chicken, duck, and quail cells. Chicken **(A)**, quail **(B)**, and duck **(C)** cells were infected with PR8 at an MOI of 5. Cells were collected for RNA-sequencing 6h post infection. Genes with a fold change 2 2 and an adjusted p-value ≤ 0.05 were considered PR8-stimulated genes (PSGs). N = 5 samples per group.

**Table S1. sgRNA sequences used in screens**

**Table S2. sgRNA sequences**

**Table S3. siRNA sequences**

**Table S4. Primer sequences**

